# A novel time-stamp mechanism transforms egocentric encounters into an allocentric spatial representation

**DOI:** 10.1101/285494

**Authors:** Avner Wallach, Erik Harvey-Girard, James Jaeyoon Jun, André Longtin, Leonard Maler

## Abstract

Learning the spatial organization of the environment is essential for most animals’ survival. This often requires the animal to derive allocentric information about the environment from egocentric sensory and motor experience. The neural circuits and mechanisms underlying this transformation are currently unknown. We addressed this problem in electric fish, which can precisely navigate in complete darkness and whose requisite brain circuitry is relatively simple. We conducted the first neural recordings in the *preglomerular complex*, the thalamic region exclusively connecting the *optic tectum* with the spatial learning circuits in the *dorsolateral pallium*. While tectal egocentric information was eliminated in preglomerular neurons, the time-intervals between object encounters were precisely encoded. We show that this highly-reliable temporal information, combined with a speed signal, can permit accurate path-integration that then enables computing allocentric spatial relations. Our results suggest that similar mechanisms are involved in spatial learning via sequential encounters in all vertebrates.

## Introduction

Learning to navigate within the spatial organization of different habitats is essential for animals’ survival (Geva-Sagiv, Las, Yovel, & Ulanovsky, 2015). Electric fish, for example, occupied a lucrative ecological niche by evolving the ability to navigate and localize food sources in complete darkness using short-range electrosensation (Jun et al., 2016). The spatial acuity that they exhibit, along with their reliance on learned landmark positions, strongly suggest that they memorize the relative arrangement of the landmarks and the environmental borders. The information animals use to generate such *allocentric* knowledge include sensory experiences collected during object encounters (Jun et al., 2016; Petreanu et al., 2012; Save, Cressant, Thinus-Blanc, & Poucet, 1998) and motor actions (heading changes and distance traveled) executed between such encounters; utilization of these motor variables in spatial learning and navigation is termed *path integration* (Collett & Graham, 2004; Etienne & Jeffery, 2004). This acquired information, however, is always *egocentric* in nature. Fittingly, the primary brain regions dedicated to sensory and motor processing, such as the *optic tectum* (OT) of all vertebrates and many cortical regions in mammals are topographically organized along an egocentric coordinate system (Knudsen, 1982; Sparks & Nelson, 1987; Stein, 1992). The neural mechanisms underlying the transformation of the egocentric sensory and motor information streams into an allocentric representation of the environment are completely unknown in any animal species; past research has not shown nor hypothesized as to which neural structures and processes are involved.

Teleost fish offer an attractive model for studying this question, as their related brain circuitry is relatively tractable: lesion studies point to the dorsolateral pallium (DL) as the key telencephalic region involved in spatial learning (Rodriguez et al., 2002), similarly to the medial cortex in reptiles and the hippocampus in mammals (see Discussion). Importantly, DL receives all sensory and motor information from OT (Bastian, 1982) via a single structure – the diencephalic preglomerular complex (Giassi, Duarte, Ellis, & Maler, 2012) (PG, **Figure 1A**); moreover, PG receives very little feedback from areas associated with DL – essentially making PG a feed-forward information bottleneck between egocentric and allocentric spatial representations. This architecture provides unique access to study egocentric-to-allocentric transformations in the vertebrate brain. In what follows we describe the first electrophysiological recordings conducted in PG of any fish species. PG has been considered part of the fish thalamus, based on simple anatomical criteria (Giassi et al., 2012; Ishikawa et al., 2007; Mueller, 2012). Our recordings provided an additional functional correspondence in that PG cells emit rapid spike bursts likely mediated by the T-type Ca^2+^ channels (**Figure 1B,C** and **Figure 1–figure supplement 1**) characteristic of the OT targets in thalamic regions of other vertebrates (Ramcharan, Gnadt, & Sherman, 2005; Reches & Gutfreund, 2009). In this contribution, we show a radical conversion of the topographic spatial representation in OT into a reliable non-topographic temporal representation of encounter sequences. We then use a computational model to demonstrate that this temporal information is readily accessible for decoding and that it is sufficiently accurate to account for spatial precision during naturalistic behavior (Jun et al., 2016).

**Figure 1:**
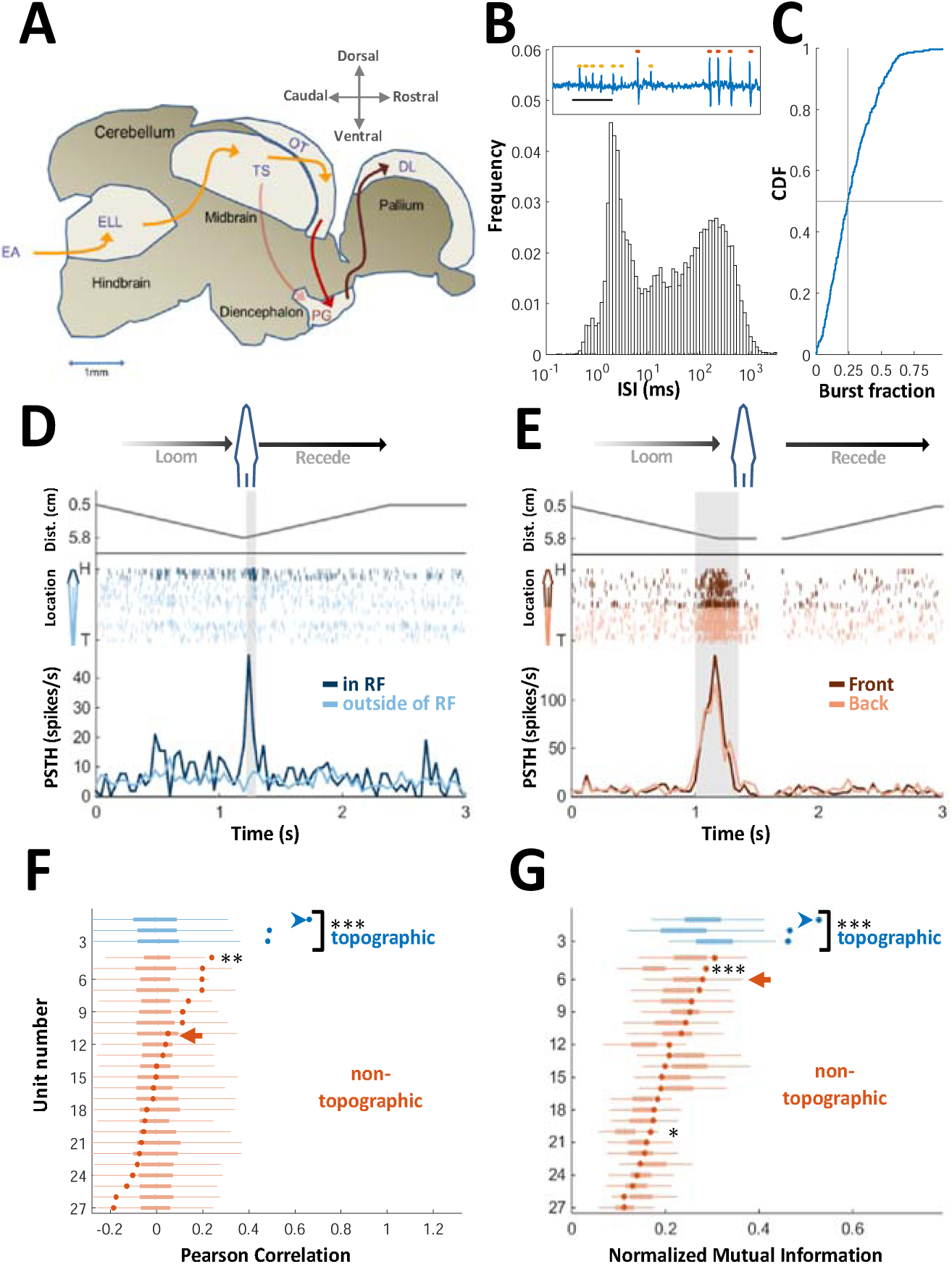
PG cells lack egocentric spatial information. **(A)** Electrosensory pathways from periphery to telencephalon. EA, Electrosensory afferents; ELL, electrosensory lobe; TS, Torus semicircularis (similar to the inferior colliculus); OT, optic tectum (homolog of the superior colliculus); PG, preglomerular complex; DL, dorsolateral pallium. **(B)** Interspike-interval (ISI) distribution in an example PG cell. Note peak around 2-ms due to thalamic-like bursting. Inset: extracellular voltage example depicting two units (red and yellow markers); scale-bar, 5 ms. **(c)** Cumulative distribution function of burst fraction (see Methods) for all PG single-units; median=0.24. **(D)** Example of an ‘atypical’ topographic PG cell with a restricted receptive field (RF) at the head. Looming-receding motions were performed at different locations. Top: object distance from skin vs. time; middle: spike raster plot, arranged according to object location (H=head, T=tail); bottom: PSTH within the RF (dark blue) and outside of it (light blue). (**E)** Example of a ‘typical’ non-topographic PG cell, responding to stimulation across the entire body. Bottom: PSTH of front (dark red) and rear (light red) halves of fish. RF extended beyond the sampled range. **(F)** Circles: Pearson correlation between normalized egocentric location and firing-rate for all units, in ascending order; box-plots: distribution of correlations obtained by random permutations of locations. Only one non-topographic cell had correlations that statistically differed significantly from those in the randomized boxes **(G)** Circles: normalized mutual information (MI) between egocentric location and firing-rate for all units, in ascending order; box plots, distribution of MI obtained by random permutations of locations. Only two non-topographic cells had MI that statistically differed significantly from those in the randomized boxes (* and ***). Blue arrowheads in (F) and (G) mark the cell depicted in (D); Red arrows mark the cell depicted in (E). P values obtained using permutation test: ‘*’, *P* < 0.05; ‘**’, *P* < 0.01; ‘***’, *P* < 0.001.

## Results

We characterized the responses of 84 electrosensory PG cells of the weakly electric fish *Apteronotus leptorhynchus* to moving objects; several motion protocols were used and the sample size for each protocol is mentioned in context. Neuronal activity was recorded extracellularly (**Figure 1–figure supplement 2**) in immobilized fish while objects (brass or plastic spheres) were moved relative to the skin using a linear motor (Methods). Cells responding to this stimulation were predominantly found in the lateral nucleus of the PG complex, PGl (**Figure 1–figure supplement 3**).

### Egocentric spatial information is abolished in PG

We first examined the spatial representation in PG cells by measuring their receptive fields (RFs; measured in 27 cells). The OT electrosensory cells driving PG have spatially-restricted, topographically organized RFs (Bastian, 1982), and thus provide labeled-line information on the egocentric position of objects. In PG, by contrast, only 11% of the cells (3/27) were topographic with a spatially restricted RF (**Figure 1D**); the majority of PG cells (89%, 24/27) responded across most or all of the fish’s body (**Figure 1E**). Therefore, PG activity does not convey a topographic ‘labeled-line’ code of object position. We also checked whether object location is encoded by the firing-rate of PG neurons (i.e., a rate code). The firing-rate was significantly correlated with object position only in one non-topographic cell (4%, 1/24 neurons; *P*<0.05, random-permutations test, **Figure 1F**). Similarly, mutual information between object position and firing-rate was significant only in two non-topographic cells (8%, 2/24 neurons; *P*<0.05, random-permutations test, **Figure 1G**). Therefore, almost all PG cells have whole-body RFs and lack egocentric spatial information– the hallmark of all electrosensory regions from the sensory periphery up to OT.

### PG cells respond to object encounters

Next, we checked what information PG neurons convey about object motion. OT cells, which drive PG, respond to an object moved parallel to the fish (longitudinal motion) while the object traverses their RFs (Bastian, 1982). Only a minority of recorded PG units (25%, 7 out of 28 tested with longitudinal motion) responded in this manner (**Figure 2A**). Rather, the majority of PG units (78%, 21/28) exhibited a strikingly different behavior, emitting a brief burst response confined to the onset (and sometimes to the offset) of object motion, but not during motion itself (**Figure 2B**). This is further demonstrated when motion in each direction was broken into four segments separated by wait periods (**Figure 2C**), evoking responses at the onset (yellow arrowheads) and offset (red arrows) of each segment across the entire body. We next applied transverse motion, which mimics an object looming/receding (incoming/outgoing) into/from the electrosensory receptive field (tested in 40 cells). Three types of responses to such motion were identified: proximity detection, encounter detection, and motion-change detection; 27.5% (11/40) displayed more than one type of response. Proximity detectors (70%, 28/40) responded when an object was encountered very close to the skin (<1.5 cm, **Figure 2D**); encounter detectors (20%, 8/40) responded when an object either entered to or departed from their electroreceptive range (∼4 cm, **Figure 2E**); lastly, motion-change detectors (47.5%, 19/40) displayed a response similar to that observed in longitudinal motion, firing at the onset/offset of motion (i.e., when the object accelerated/decelerated, **Figure 2F**); remarkably, this type of response was relatively distance-invariant, yielding comparable responses both very close to (0.5 cm) and very far from (5.5 cm) the skin despite the drastic effects of distance on both the magnitude and spread of the object’s electrical image (**Figure 2–figure supplement 1**, see Chen, House, Krahe, & Nelson, 2005).

**Figure 2:**
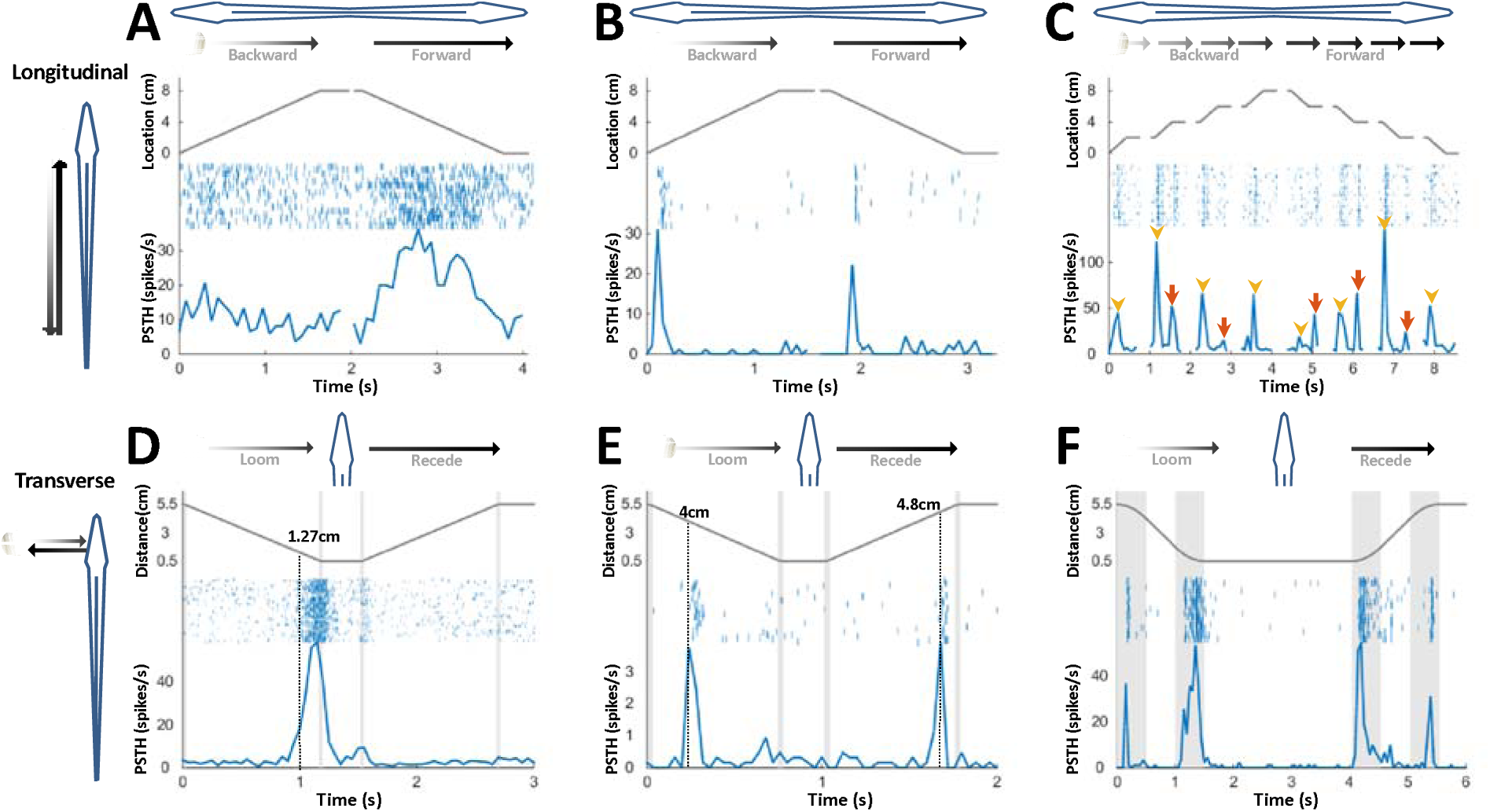
PG cells respond to motion novelty. **(A-C)** Longitudinal motion. Top: object location (distance from the tip of the nose); middle: spike raster; bottom: PSTH. **(A)** ‘Atypical’ PG cell responding uni-directionally, throughout the motion toward the head. **(B)** ‘Typicial’ PG cell responding to motion-onset at the head during backward motion and to motion-onset at the tail during forward motion. **(C)** PG cell responding to motion onset (yellow arrowheads) and to motion offset (red arrows) in intermittent-motion protocol. **(D-F)** Responses to transverse (looming-receding) motion. All PG cells recorded with transverse-motion (40/40) responded to object looming, while only 40% (16/40) responded to receding. Top: object distance from skin; middle: spike raster; bottom: PSTH. Grey shadings mark acceleration/deceleration. **(D)** Proximity detector cell. Note that the response starts prior to the deceleration (shaded). **(E)** Encounter detector cell. **(F)** Motion change detector cell, responding when objects accelerated/decelerated (shaded, 10 cm/s^2^ in this example) similarly to the response observed to longitudinal motion (panels B and C).

We conclude that PG electrosensory cells predominantly respond to novelty: onset/offset of object motion within their receptive field or the introduction/removal of objects into/from this receptive field; we use the term object encounters to designate all such events. Most PG cells responded both to conductive and non-conductive objects (**Figure 2–figure supplement 2**) and would therefore not discriminate between different object types, e.g., plants versus rocks. We can, therefore, infer that during active exploration of the environment, PG reports to the dorsal pallium whenever the fish encounters or leaves a prey or a landmark (e.g., root mass or rock) or alters its swimming trajectory near a landmark.

### PG cells display history-independent, spatially non-specific adaptation

While egocentric spatial information was scarce in PG, temporal information was prevalent. Many of the PG cells exhibited pronounced adaptation to repeated motion (45%, 15/33 neurons tested with repeated-motion protocol; **Figure 3–figure supplement 1**). We delivered a sequence of object encounters (motions in and out of the RF, **Figure 3A**); time-intervals between sequential encounters were drawn randomly and independently (in the range of 1–30 s). The response intensity (number of spikes within an individually determined time window) of these adapting cells to each encounter was strongly correlated with the time-interval immediately prior to that encounter (**Figure 3B**). By contrast, there was no significant correlation with intervals prior to the last one (inset of **Figure 3B** and **Figure 3C**). Thus, a large subset of PG cells encoded the duration of the time-interval *preceding the last encounter* – but did not convey information about the intervals prior to that. We term this history-independent adaptation. To quantify this insensitivity to prior history, the response of each cell was modeled using a use-dependent resource variable with memory parameter f]; the memory of past intervals is more dominant as f] → 1, while f] = 0 for a completely history-independent cell (**Figure 3–figure supplement 2**). We found that 47% of the adapting cells (7/15) were best fitted with f] < 0.1, and only 13% (2/15) had f] > 0.5 – meaning that most cells indeed carried little information about time-intervals preceding the last one. Interestingly, similar behavior was also found in visually-responsive PG cells (**Figure 3–figure supplement 3**). This result contrasts with many studies in the mammalian cortex, where adaptation is incremental and responses depend on multiple preceding intervals (Lampl & Katz, 2017; Ulanovsky, Las, Farkas, & Nelken, 2004). Moreover, a spatial ‘oddball’ experiment (Methods), which we conducted on 14 neurons (6 of which were adapting) revealed that this adaptation is also spatially non-specific – i.e. encounters at one body location adapt the response to encounters at all other locations (**Figure 3D-F**). We thus discovered here an entirely novel adaptation mechanism in the vertebrate thalamus that enables neurons to encode the time between the two most recent successive stimuli, without contamination by prior encounter history and irrespective of the objects’ egocentric position.

**Figure 3:**
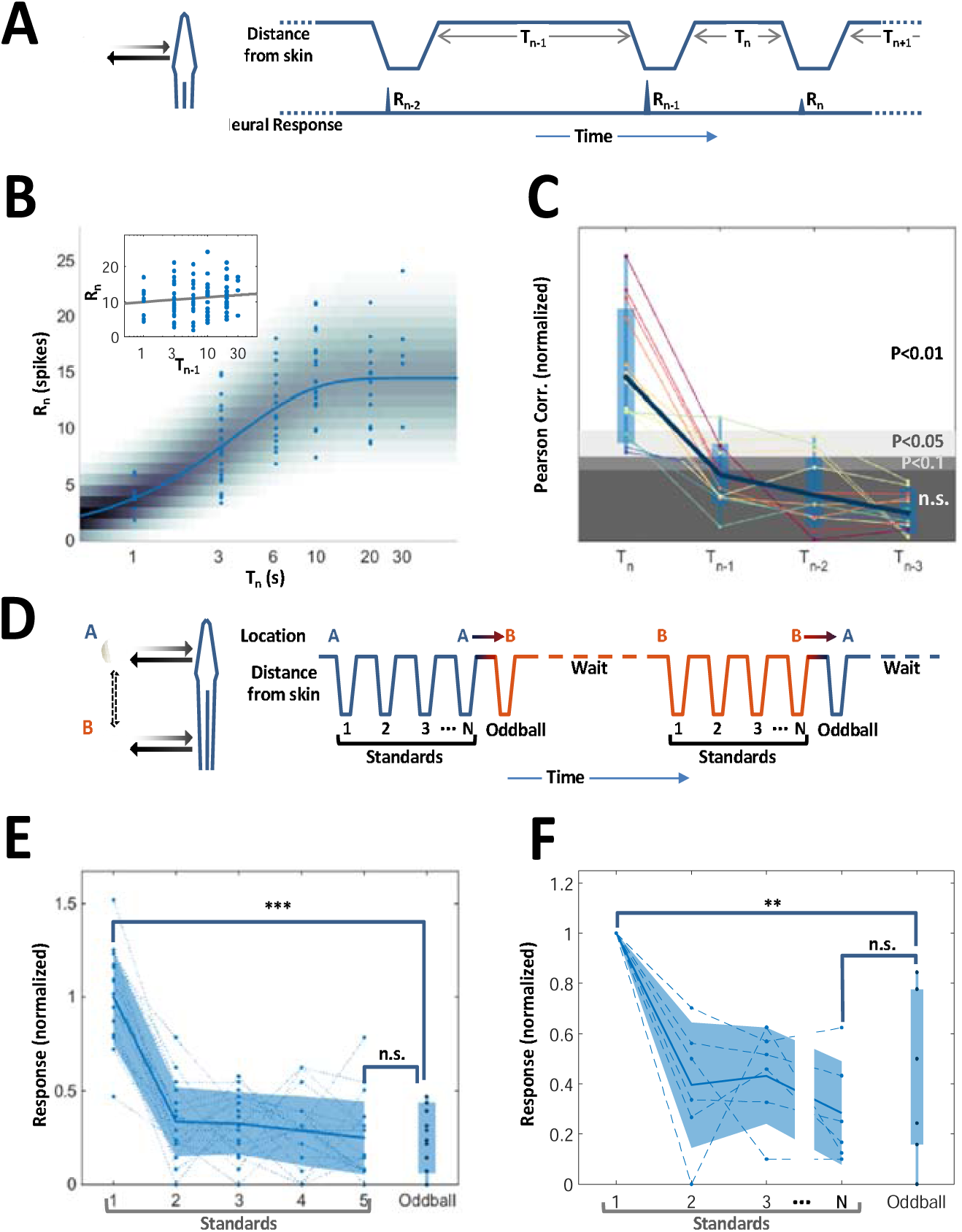
History-independent, spatially non-specific adaptation of PG cells. **(A)** Random interval protocol. Object motions toward- and away-from-skin generated responses {R_l_}, measured by spike count within individually determined time-window. Motions were interleaved by random time intervals{T_l_} (1–30 s). **(B)** Response versus the preceding time-interval (log scale) for an example neuron (circles). Model fitted to the data-points is depicted by response rate (solid blue line) and spike count distribution (background shading); correlation of data to model=0.56 (P < 10^−6^, random permutations). Inset: same data-points plotted as function of penultimate intervals; grey line: linear regression to log(T) (correlation=0.15, P = 0.136, random permutations). **(C)** Correlations between responses and time-intervals (last interval and one-, two- or three-before-last), normalized to the 10, 5 and 1 top precentiles of correlations generated by random permutations of the data, for each of the 14 adapting cells tested (thin colored lines; thick blue line: mean across population). For most cells, the response was significantly correlated only with the last interval. **(D)** Spatial oddball protocol. A sequence of N standard stimuli at one location (N=5-9) was followed by one oddball stimulus at a different location. **(E)** Response of example cell (normalized to non-adapted response) to oddball protocol (dotted lines: individual tirals, N=10; thick line and shaded area: mean±standard deviation). Response to the oddball was significantly weaker than the non-adapted response (P < 10^−4^, bootstrap) and not significantly different from the response to the last standard stimulus (P = 0.45, bootstrap). **(F)** Normalized response of all adapting cells tested with the spatial oddball protocol (dashed lines: individual cells, N=6; thick line and shaded area: mean±standard deviation). Population response to the oddball was significantly weaker than the non-adapted response (P =0.001, bootstrap) and not significantly different from response to the last standard stimulus (P = 0.2, bootstrap). ‘**’, P < 0.01; ‘***’, P < 0.001; n.s., not significant.

### PG activity explains path integration acuity

Can PG activity be used to explain observed spatial behavior in freely-swimming electric fish? PG is the only source of sensory information for the dorsolateral pallium (DL), the likely site of spatial memory (Rodriguez et al., 2002). Therefore, we hypothesized that the history-independent adapting PG cells provide DL with one *necessary* component for path integration: the temporal sequence of object encounters (the two other components are heading-direction and linear velocity). To demonstrate this, we constructed a bootstrap model of the PG population (Methods). This model was then stimulated at random time-intervals. The last time-interval could be precisely decoded from the population response using an unbiased Maximum-Likelihood Estimator (MLE, **Figure 4A**, Methods). As expected, the estimation error increased with the time-interval being decoded and decreased with population size (**Figure 4B**). Given the animal’s velocity, the model’s estimated interval may be converted to traveled distance (**Figure 4C**). Finally, we compared the model’s estimation error with the behavioral errors exhibited by fish trained to find food at a specific location, either directly after leaving the tank boundary (long distance to food) or after leaving a memorized landmark (shorter distance to food) (Jun et al., 2016) (**Figure 4–figure supplement 1**). The distributions of the temporal and spatial behavioral errors (boxplots in **Figure 4B,C**), both with and without landmarks, were captured by MLE using only ∼500 adapting cells. The lateral subdivision of PG (PGl), in which most motion-responsive cells were found, contains about 60,000 cells (Trinh, Harvey-Girard, Teixeira, & Maler, 2015); based on our data we can estimate that around 21% of these are history-independent adapting cells (7/33), i.e. close to 13,000. Thus, the number of PGl cells is *sufficient* to attain the observed behavioral precision, even when additional encoding errors (e.g. in heading and velocity estimation) are taken into account. Taken together, our simulations suggest that time-interval encoding in PG can consistently account for the observed behavioral precision of spatial learning.

**Figure 4:**
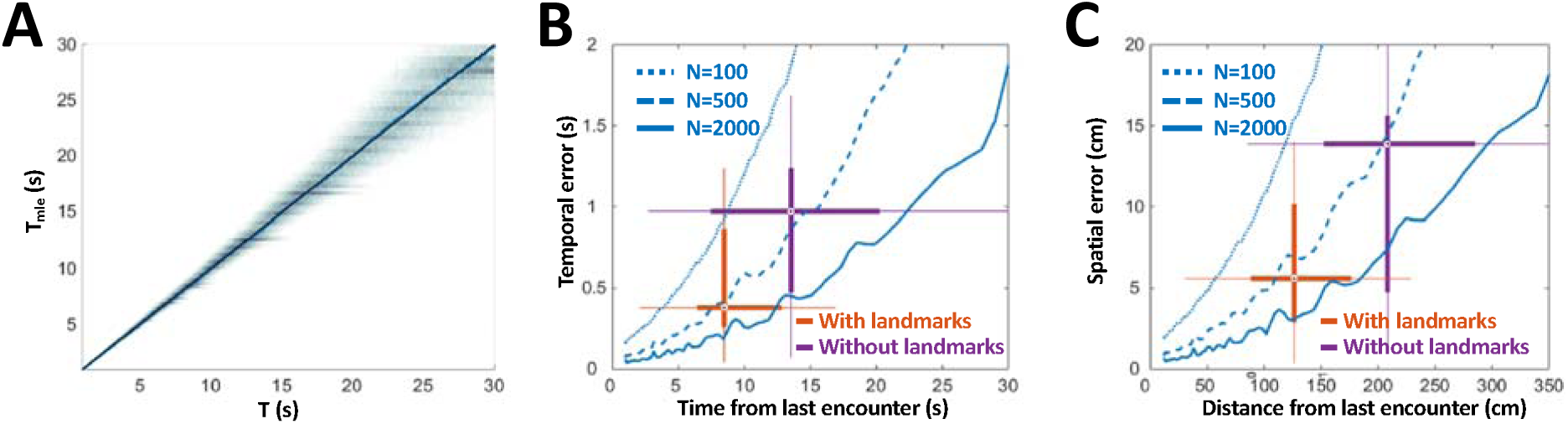
PG simulation explains the accuracy of animals’ spatial behavior. **(A)** Distribution of MLE of time interval vs. actual time interval (using *N* = 500 cells). Estimation is unbiased: the mean (blue line) follows the identity diagonal. **(B)** Temporal representation. Blue curves: mean MLE error with 100 (dotted), 500 (dashed) and 2000 cells (solid). Two-dimensional boxplots: distribution of behavioral temporal estimation errors vs. time from last encounter (red: with landmarks, purple: without landmarks, see Methods and **Figure 4–figure supplement 1**). **(C)** Spatial representation. MLE temporal results were converted to distance by multiplying by the median velocity reported for navigating fish (12 cm/s) (Jun, Longtin, & Maler, 2016).

## Discussion

Our results suggest the following egocentric-to-allocentric transformation scheme: PG derives a sequence of discrete novelty events (encounters) from OT activity. The remarkable history-independent adaptation process provides an accessible, accurate and unbiased representation of the time intervals between encounters. We have found visually-responsive PG cells displaying similar adaptation features (**Figure 3–figure supplement 3**), suggesting that this mechanism is implemented across multiple sensory modalities. The elimination of egocentric spatial information in PG – both in the response itself and in its adaptation – ensures that the encoding of time is invariant to the specific body part encountering the object. This temporal information is then transmitted to DL, which can use it to integrate the fish’s swim velocity to obtain distance-traveled, a key allocentric variable. For fish, the necessary velocity information may be provided by the lateral-line system (Chagnaud, Bleckmann, & Hofmann, 2007; Oteiza, Odstrcil, Lauder, Portugues, & Engert, 2017); several lateral-line responsive PG units were encountered in our recordings (**Figure 4–figure supplement 2**). Finally, DL can combine the distance information with instantaneous heading-direction (vestibular system, Straka & Baker, 2013; Yoder & Taube, 2014) to yield the animal’s allocentric spatial position (Etienne & Jeffery, 2004). A recent study in zebrafish suggests that DL neurons can indeed process temporal information on the long time-scales discussed here (Cheng, Jesuthasan, & Penney, 2014). Our computational analysis demonstrates that the PG temporal information is sufficient to account for the spatial acuity displayed in behavioral studies of gymnotiform fish utilizing electrosensory information alone (**Figure 4 Figure 4–figure supplement 1**).

Our findings shed light on the primitive basis of egocentric-to-allocentric transformations. Short-range sensing, used by ancestor species living in low-visibility environments, necessitated the perception of space through temporal sequences of object encounters. With the evolution of long-range sensory systems such as diurnal vision (MacIver, Schmitz, Mugan, Murphey, & Mobley, 2017), simultaneous apprehension of the spatial relations of environmental features became possible. The neural mechanisms implementing sequential (space-to-time) spatial inference and simultaneous spatial inference presumably both exist in mammals, e.g. we can acquire a map of relative object locations by looking at a scene from afar, or by walking and sequentially encountering landmarks with our eyes closed. Whether the sequential or the simultaneous spatial-inference is more dominant may depend on the species (e.g., nocturnal or diurnal) and on context (e.g., open field or underground burrows). However, it is not clear whether sequentially versus simultaneously-acquired spatial knowledge is processed in a common circuit. Indeed, clinical case studies on the regaining of eyesight in blind humans indicate that sequential and simultaneous spatial perceptions are fundamentally different, and may therefore involve two distinct computations and neuronal pathways (Sacks, 1995). The population of thalamic neurons that we discovered may underlie one of these two major computations – the encoding of sequential temporal information – and we hypothesize that such neurons underlie sequential spatial learning in all vertebrates.

There is substantial evidence indicating that the pathway studied here indeed has parallels in other vertebrates, and specifically in mammals. PG’s homology to posterior thalamic nuclei is supported by previously published anatomical and developmental findings (Giassi et al., 2012; Ishikawa et al., 2007; Mueller, 2012), as well as by physiological (**Figure 1B,C**) and molecular (**Figure 1–figure supplement 1**) results presented here. The thalamic pulvinar nucleus is particularly similar to PG in that it receives direct tectal input (Berman & Wurtz, 2011). Its involvement in visual attention and saliency in primates (Robinson & Petersen, 1992) corresponds to PG’s involvement in novelty detection (**Figure 2**). Moreover, pulvinar lesions are associated with saccadic abnormalities and deficits in the perception of complex visual scenes (Arend, Rafal, & Ward, 2008), suggesting a link to the sequential mode of spatial learning. Komura et al. (2001) demonstrated that posterior thalamic regions (including the rodent equivalent of the pulvinar) can implement interval timing computations over long time-scales (>>1s); however, the mechanistic basis for these computations has not been identified (Simen, Balci, de Souza, Cohen, & Holmes, 2011) and potential contributions to path integration have not been explicated. Finally, a recent study in rodents demonstrated spatially non-specific adaptation in VPM (Jubran, Mohar, & Lampl, 2016), a posterior thalamic nucleus responding to vibrissal object encounters (Yu, Derdikman, Haidarliu, & Ahissar, 2006). Taken together, we hypothesize that thalamic space-to-time mechanisms akin to those presented here play an important role in mammalian sequential spatial learning, especially in nocturnal animals relying on sparse sensory cues (Save et al., 1998).

The telencephalic target of PG, DL, resembles the mammalian hippocampus not only in function, as revealed in lesion studies (Rodriguez et al., 2002), but also in development, gross circuitry and gene expression (Elliott, Harvey-Girard, Giassi, & Maler, 2017). The role of the hippocampus in spatial learning and navigation is well established, and hippocampal neural correlates of allocentric spatial variables have been exquisitely described (Barry & Burgess, 2014; Buzsaki & Moser, 2013). There is also evidence for the importance of time coding in the mammalian hippocampus: ‘Time cells’ responsive to elapsed time have been reported and, in some cases, these cells also respond at specific spatial loci (Eichenbaum, 2014). Furthermore, a recent study on the representation of goals in the hippocampus found cells encoding the length/duration of the traveled path (Sarel, Finkelstein, Las, & Ulanovsky, 2017). The mechanism we have found may therefore contribute to creating temporal coding in the hippocampus, not just in the context of egocentric-to-allocentric transformations but rather whenever expectations associated with specific time intervals need to be generated. It should be noted, however, that unlike DL’s direct thalamic input via the PG bottleneck, the hippocampus receives sensory and motor information primarily via the cortex. Furthermore, multiple bi-directional pathways connect the mammalian sensory and motor cortical regions with the hippocampal network. Pinpointing the exact loci where egocentric-to-allocentric transformations may take place in the mammalian brain is therefore extremely challenging. We propose that this transformation is initiated in the mammalian thalamus where history-independent adaptation also encodes time between encounter events. Finally, we propose that this thalamic output contributes to the generation of an allocentric spatial representation in the mammalian hippocampus.

## Author Contributions

A.W. performed the electrophysiological experiments, data analysis and modeling; E.H.G. performed PCR experiments; J.J.J. performed behavioral experiments; A.L. and L.M. supervised the project; A.W., A.L. and L.M. wrote the manuscript.

## Acknowledgments

We thank Bill Ellis for technical support and Nachum Ulanovsky for comments and suggestions. This research was supported by NSERC Grant 121891-2009 (to A.L.), NSERC Grant 147489-2017 (to L.M.) and Canadian Institutes of Health Research Grant 49510 (to L.M. and A.L.).

## Methods

### Experimental Model

All procedures were approved by the University of Ottawa Animal Care and follow guidelines established by the Society for Neuroscience. *Apteronotus leptorhynchus* fish (imported from natural habitats in South America) were kept at 28°C in community tanks. Fish were deeply anesthetized with 0.2% 3-aminobenzoic ethyl ester (MS-222; Sigma-Aldrich, St. Louis, MO; RRID: SCR_008988) in water just before surgery or tissue preparation.

### Surgical procedure for in-vivo recordings

Surgery was performed to expose the rostral cerebellum, lateral tectum and caudal pallium. Immediately following surgery, fish were immobilized with an injection of the paralytic pancuronium bromide (0.2% weight/volume), which has no effect on the neurogenic discharge of the electric organ that produces the fish’s electric field. The animal was then transferred into a large tank of water (27□°C; electrical conductivity between 100–150□μS□cm^−1^) and a custom holder was used to stabilize the head during recordings. All fish were monitored for signs of stress and allowed to acclimatize before commencing stimulation protocols.

### Neurophysiology

Custom made stereotrodes or tritrodes were made of 25-μm diameter Ni-Cr wire (California Fine Wires). Each electrode was manually glued to a pulled filamented glass pipette (P-1000 Micropipette Puller, Sutter Instrument, Novato, CA); the glass pipette provided mechanical rigidity that allowed advancing the tetrode to the deep-lying PG. Prior to recording, tetrode tips were gold-plated (NanoZ 1.4, Multi Channel Systems, Reutlingen, Germany) to obtain 200–300 kOhm impedance at 1kHz. The electrode was positioned above the brain according to stereotaxic brain atlas coordinates (150-300 µm caudal to T26 and 800-1000 µm lateral to midline (Maler, Sas, Johnston, & Ellis, 1991)), and lowered using a micropositioner (Model 2662, David Kopf Instruments, Tujunga, CA) while delivering visual and electrosensory stimuli. Tectal responses to such stimuli (Bastian, 1982) were usually detected twice, around 1200 µm and around 1900 µm ventral to the top of cerebellum (as expected from the curved shape of OT). The electrode usually then transversed nucleus Electrosensorius, producing weak multi-unit responses to electrocommunication stimuli around 2300-2500 µm (Heiligenberg, Keller, Metzner, & Kawasaki, 1991). PG units were usually encountered between 2800 µm and 3400 µm ventral to the top of the cerebellum, and were easily identified due to their characteristic rapid spike bursts (**Figure 1B,C**). Differential extracellular voltage was obtained by using one stereotrode/tritrode channel as reference. This enabled near-complete cancellation of the electric organ discharge (EOD) interference.

We report on the responses of 84 PG neurons responsive to object motion; several motion protocols were used and the sample size for each protocol is mentioned in context. We also found PG cells responding to electrocommunication signals, mostly within the medial subdivision of PG (PGm). As expected from the sparse retinal input to OT (Sas & Maler, 1986), we recorded only a small number (*n* = 19) of PG cells responsive to visual input (**Figure 3– figure supplement 2**). In addition, we identified a small number of cells responsive to passive electrosensory (Grewe, Kruscha, Lindner, & Benda) (ampullary receptors, *n* = 27), acoustic (*n* = 7) and lateral line (*n* = 7, **Figure 4–figure supplement 2**) stimulation – but did not attempt to further characterize their coding properties.

Cell responses were initially manually tested with brass and plastic spheres. Cells responding to both were also tested with an electrically neutral gel ball made of 15% agarose in tank water to exclude lateral-line responses (Heiligenberg, 1973) (**Figure 4–figure supplement 2**). A plastic or brass sphere (1.21 cm diameter) was connected to an electromechanical positioner (Vix 250IM drive and PROmech LP28 linear positioner, Parker Hannifin, Cleveland, OH), which was pre-programmed for the appropriate motion sequence and initiated by outputs from our data acquisition software (Spike2, Cambridge Electronic Designs, Cambridge, UK). Typically, a trapezoidal velocity profile was used with 150 cm/s^2^ acceleration and 5 cm/s peak velocity, and total distance in either direction of 8 cm (longitudinal) or 5 cm (transverse). For protocols involving motion in two axes (receptive field sampling, Oddball), the object was attached to a second electromechanical positioner (L12-100-50-12-P linear actuator, Firgelli Technologies, Ferndale, WA), which was mounted perpendicularly to the first one. In order to measure the receptive field (RF) size, we used one motor to repeatedly perform transverse object motion (towards and away from the fish) while a second motor was used to randomly change the longitudinal position between repetitions. This protocol was performed on a total of 27 cells. To check the spatial specificity of the adaptation process (i.e. if it is affecting only the body location experiencing the encounters or a whole-body effect), we performed a spatial oddball experiment: First, a series of N ‘standard’ encounters (in and out transverse object motion, N=5-9) were given in rapid succession at one location (e.g. the head); the (N+1)^th^ encounter was given at a different location (e.g. the trunk) while maintaining the same time-interval between encounters (3-5 s, the second motor was used to quickly switch positions). Each such series was followed by a long recovery period (≥30s) and then repeated in the opposite direction. This was performed on a total of 14 cells, out of which 6 were found to be adapting (in either direction).

### Histology

In the initial experiments the location of PG was verified by preparing histological sections and locating the electrode track marks (**Figure 1–figure supplement 3)**. After recordings were complete, the fish was deeply anesthetized using tricaine methanesulfonate (MS-222 0.2 g/L; Sigma-Aldrich, St-Louis, Mo) and transcardially perfused with 4% paraformaldehyde, 0.1% glutaraldehyde, and 0.2% picric acid in 0.1 M PBS pH 7.4. Brains were removed and incubated overnight in a solution of 4% paraformaldehyde, 0.2% picric acid, and 15% sucrose in 0.1 M PBS pH 7.4 at 4°C. Cryostat sections were cut at 25 μm in the transverse plane and mounted on Superfrost Plus glass slides (Fisher Scientific, Pittsburgh, PA). All sections were counterstained with green fluorescent Nissl reagent 1:300 (Molecular Probes, Eugene, OR; NeuroTrace 500/525 green-fluorescent Nissl Stain #N21480, RRID: SCR_013318) in PBS for 20 minutes at room temperature.

### Expression of T-type Ca^2+^ channel α-subunits

Three adult male fish were anesthetized. Ice-cold ACSF was dripped on the head while the skull was removed and brains quickly removed and submerged in ice-cold ACSF. PG and DL brain regions were superficially located (Maler et al.), identified, dissected out and stored at -20°C. Tissues were weighted and total RNA was purified using Trizol (Sigma-Aldrich) according to the manufacturer’s recommendations. Contaminating genomic DNA was digested with DNAse1, total RNA was then precipitated overnight at -20 °C, resuspended in nuclease-free water and quantified by spectroscopy. 300 ng of total RNA were used for first strand cDNA synthesis using the Maxima H Minus First Strand cDNA Synthesis Kit (K1681; ThermoFisher). PCR were made with the Taq polymerase (EP0402; ThermoFisher) according to manufacturer’ recommendations for 35 cycles using the following primer pairs: G amplicon (301bp), direct: 5’-CGACACCTTCCGCAAAATCG-3’, reverse: 5’-AGCACAGACAGACCTCCGc-3’; H amplicon (338bp), direct: 5’-GGGACGATTTCAGGGACAGG-3’, reverse: 5’-CACTCGCAGCAGACGGAA-3’; I amplicon (355bp), direct: 5’-TGGGATGAGATTGGAG TGAAAC-3’, reverse: 5’-AGCGGACCAGCTTAATGACC-3’. Amplicons were then migrated on a 1% agarose Et-Br gel and photographed with a BIO-RAD Gel Doc System.

### Quantification and statistical analysis

#### Preprocessing

Data were acquired using Spike2 (Cambridge Electronic Designs, Cambridge, UK) and analyzed in Matlab (MathWorks, Natick, MA). Single units were sorted offline by performing principal component and clustering analyses. Units were considered to be well-separated and to represent individual neurons, only if (a) their spike shapes were homogenous over time and did not overlap with other units or noise; and (b) the unit exhibited refractory period of >1 ms in autocorrelation histograms (**Figure 1–figure supplement 2**).

#### Burst analysis

Any spike preceded or followed by an ISI ≤ 8 ms was regarded as one participating in a burst. This was justified by observing the ISI first-return maps (Ramcharan et al., 2005), which showed characteristic clusters separated around 8-ms. To compute the burst fraction, the number of spikes participating in bursts was divided by the total number of spikes emitted by the unit.

#### Information content

To determine the information the spiking response *r* contains on object position *p*, the normalized mutual information was estimated: 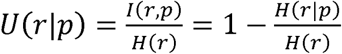, where *I* is the mutual information and *H* is the information entropy. The conditional entropy *H(r|p)* was calculated by binning the responses obtained in the RF sampling protocol (**Figure 1D,E**) into 24 bins and calculating the conditional response (spike count) distribution in each bin. Note that 0≤*U(r|p)*≤1, with *U(r|p)=0* when the spiking is completely independent of position, and *U(r|p)=1* when the firing is completely predictable given the object position.

#### Statistics

In all paired comparisons, the bootstrap method (5,000 random redistributions without replacement) was employed to estimate statistical significance. Random permutations were used to evaluate significance of correlations (5,000 random permutations without replacement). Sample sizes were determined by statistical requirements, aiming at confidence levels > 95%. No statistical methods were used to pre-determine sample sizes but our sample sizes are similar to those generally employed in the field. No randomization or blinding was used is this study.

#### Modeling and Simulations

##### Dynamic adaptation model

The random interval protocol (**Figure 3A**) produced for each cell an interval (input) vector {*T*_*n*_} and a spike count response (output) vector {*R*_*n*_} (count computed for each unit in a time-window determined by its response type, see **Figure 2**). For some of the cells, a slow decline in response due to experimental instability was corrected by dividing the response time course by a least-squared fitted slow (>10 min) exponential decay. The model (**Figure 3–figure supplement 2)** with dynamics following each encounter givenby

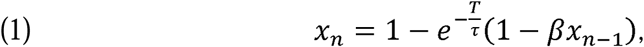

where *T*_*n*_ is the last time interval, β is the memory coefficient (0“’”1 and, the recovery (Tsodyks & Markram, 1997), where ’.1-,; we chose this formalization so that the extent of history-dependence in the adaptation dynamics will be emphasized (i.e., ’.0 signifies complete history independence and history dependence increases as β →1)

The neuron’s firing parameter λ at the n^th^ encounter is λ *=*, where *a>0, c∈*and squares, while β and τ, were found by exhaustive search on a 2-D grid. Fitting was deemed denotes non-negative thresholding. Fitting of *a* and *c* was performed using linear least successful if the parameter values generated by the model were correlated with the actual responses with *P* < 0.05 (random permutations). This was true for 15/33 cells. Note that when β =0 (‘history-independent’ adaptation), we get:

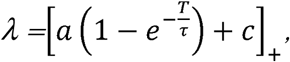

which corresponds to the solid blue line in **Figure 3B**.

Finally, {*λ*_*n*_} were used as the parameters of Poisson random variables to generate stochastic spike-counts *{R*_*n*_*}*, so that the number of spikes emitted at each encounter was distributed according to:

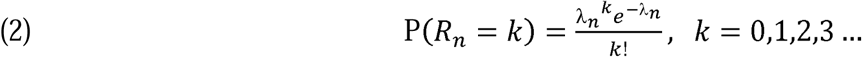

##### Simulation of bootstrap population

The set of parameter values obtained for all 15 fitted adapting cells (a, c and τ) were randomly drawn from to generate a bootstrap population of N cells (N=100, 500 and 2000). The memory variable β was set to 0 for all cells to simplify decoding. We also assumed that spiking across the population was statistically independent with Poisson statistics given the last interval, i.e.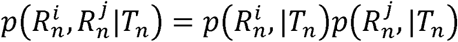 where 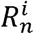 is the response of neuron *i* to time interval *n*. Random intervals were drawn in the range 1–30 s, and for each interval a vector of responses (spike counts) across the population 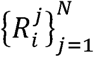 was generated, where N is the population size.

##### Maximum-Likelihood Estimation

Since 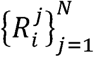 are independent Poisson r.v.’s, with each r.v. distributed according to equation (2), the likelihood of this population response is and the log-likelihood is time interval

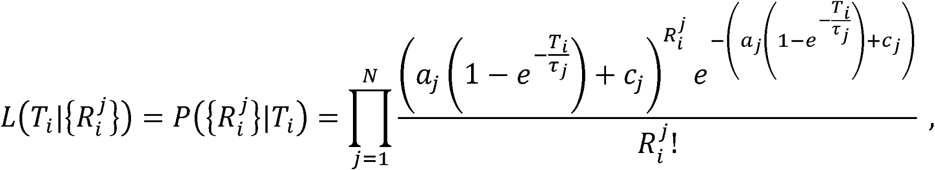

and the log-likelihood is

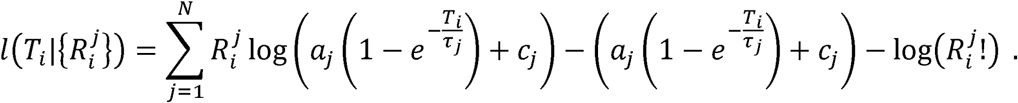

The Maximum-Likelihood Estimator of the last time interval is therefore obtained by finding the that maximizes this likelihood:

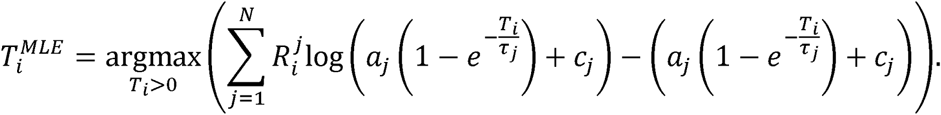

This maximum was found numerically for each generated time interval.

### Analysis of behavioral data

Published spatial learning data (Jun et al., 2016) were used; methods used for the generation of these data are explained in detail in that study. Briefly, South American weakly electric fish (*Gymnotus* sp.) were trained to find food (a mealworm restrained to a suction cup) in complete darkness, at a specific location within a 150 cm diameter custom made circular arena. In experiments with landmarks, four acrylic objects (two square prisms, 5.6 and 9.0 cm/side and two cylinders, 7.6 cm and 10.2 cm diameter) were placed in fixed locations within the arena. In each daily session, each fish was given four trials to find the food. After a 12-session training stage, animal performance stabilized, and four test sessions were performed in which food was omitted in one randomly assigned ‘probe’ trial. Only data from these four last sessions were analyzed here. A total of four fish were used with landmarks and eight fish were used without landmarks. Fish behavior was video recorded and tracked as previously described (Jun, Longtin, & Maler, 2014).

The cruising time/distance from the last encounter (with either a landmark or the tank wall) to the location of the food is the epoch/trajectory the fish had to memorize in order to perform path integration. This was measured in each of the ‘food’ trials (**Figure 4–figure supplement 1A,B**); the segment from the last encounter (3 cm from a landmark or 6 cm from the arena wall) to the detection of food (3 cm from food location) was found and the trajectory total length and duration were computed. The spatial ‘decoding’ errors were obtained by measuring where the fish searched for the missing food in the probe trials (**Figure 4–figure supplement 1C,D**); the normalized histogram of the visiting frequency across space (heat map) was fitted with a two-dimensional Gaussian function described by six parameters (μ_x_, μ_y_, σ_x_, σ_y_, θ, A: colored oval contours). The distance between the Gaussian center (μ_x_,μ_y_) (white ‘x’ mark) and the food location (white ‘+’ mark) is the spatial error; dividing this error by the median velocity in the trial produces the temporal error.

## Figure Supplements

**Figure 1–figure supplement 1.**
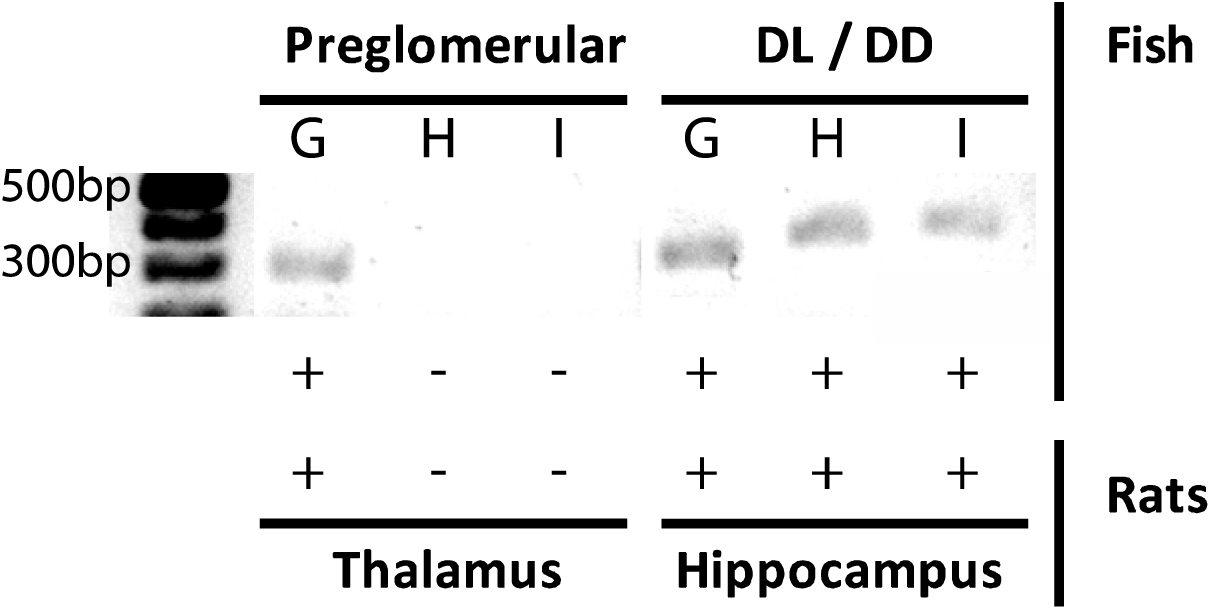
T-type Ca^2+^ channel expression profiles support PG thalamic homology. mRNA expression of G, H and I subtypes of the T-type Ca^2+^ channel α-unit, for PG (left) and the pallial DL-DD region (right) of *A. leptorhynchus* – compared with mRNA expression of the same subunits in the thalamus and hippocampus of rats (Talley et al., 1999). Only the α1G transcript of the T type Ca^2+^ channel (Ca_V_T) is expressed in both PG and thalamic neurons; in contrast, all three transcripts are expressed in DL and hippocampal neurons. We propose that PG spike bursting (**Figure 1B,C**) is generated by the α1G T-type Ca^2+^ channels and that this is a highly conserved biophysical/genetic feature of neurons in the vertebrate thalamus.

**Figure 1–figure supplement 2.**
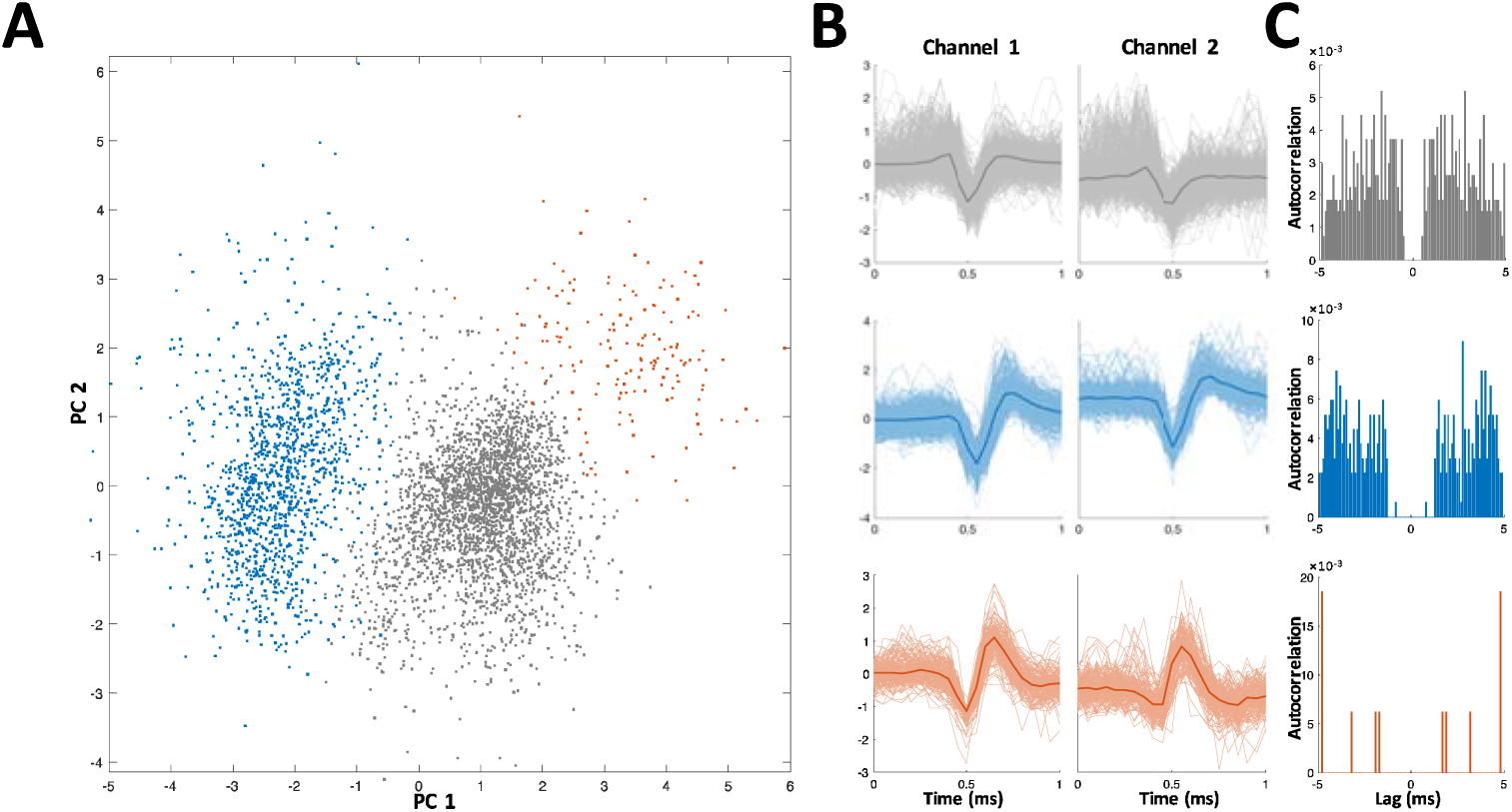
Spike sorting example. Recordings were made using either stereotrodes (two wire bundles) or tritrodes (three wire bundles); one wire was used as reference, so that near perfect cancelation of the electric organ discharge (EOD) interference was obtained. Recordings therefore consisted of either one or two channels. In the depicted example two channels (i.e., a tritrode) were employed. (**A**) Distribution of first two principal components of detected spikes, color coded according to unit identity (grey-multi-unit, blue and red-single units). (**B**) Spike shapes in the two recording channels for each unit (color codes as in panel A). (**C**) Autocorrelations of spiking activity in the three units. Single units (blue and red) are characterized by a refractory period (larger than the refractory period set for the spike detection algorithm, 1 ms).

**Figure 1–figure supplement 3.**
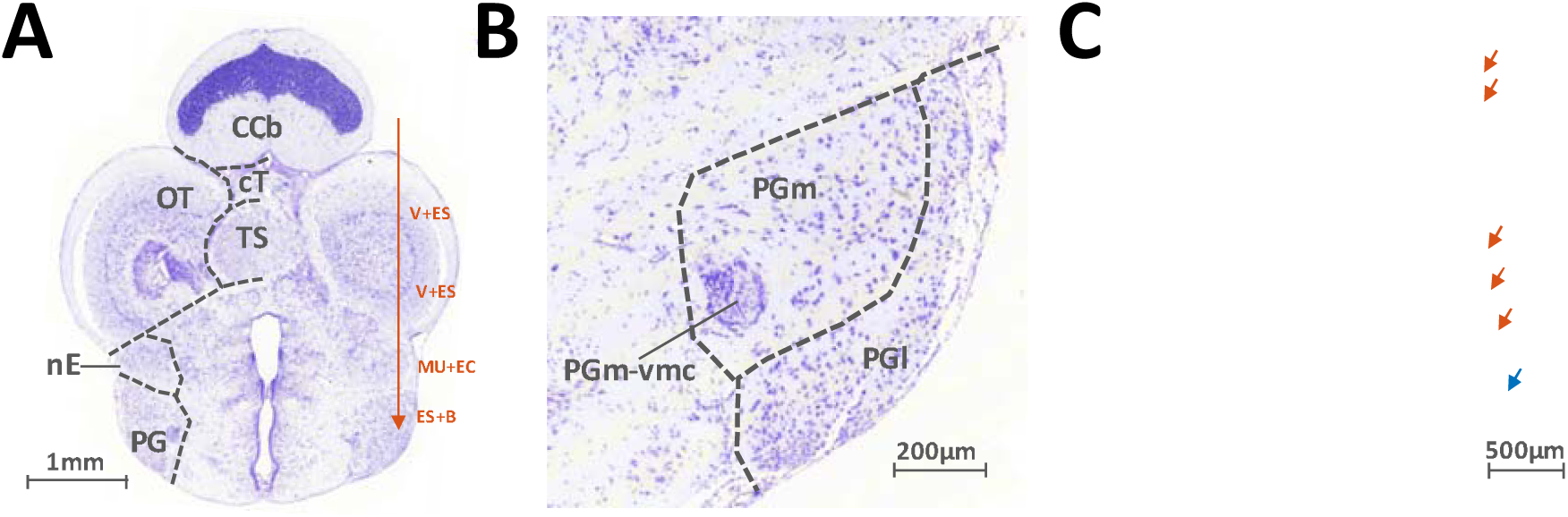
Histology. (**A**) Transverse slice of the brain of *A. Leptorhynchus*, about 300 μm caudal to the midbrain-telencephalon boundary (around electric fish atlas level T24, Maler et al., 1991). Electrodes were inserted 800-1000 μm lateral to midline and advanced ventrally (red arrow). Visual and electrosensory stimuli were presented as the electrode was lowered in order to identify PG. There was variation in the depth of the various responses encountered depending on the size and precise orientation of the fish. Typically, visual and electrosensory (V+ES) responses were encountered around 1200 μm and 1900 μm ventrally to the top of the corpus cerebellum (CCb), as the electrode passed through the rostral edge of the optic tectum (OT, Bastian, 1982). Further down, weak multi-unit activity responding to electro-communication signals (MU+EC) was usually detected around 2300-2500 μm, as the electrode passed through n. electrosensorius (nE, Heiligenberg et al., 1991). Finally, preglomerular (PG) responses were predominantly electrosensory and were characterized by bursting (ES+B, see Figure 1B-C). TS: Torus Semicircularis; cT: tectal commissure. **(B)** Close-up image of panel (A), showing the lateral (PGl) and medial (PGm) subdivisions of PG. PGm-vmc: ventromedial cell group of PGm. **(C)** Transverse slice prepared after recording motion-sensitive PG cells. The electrode track is visible through the edge of the cerebellum, the bottom of OT and nE (red arrows). The electrode tip mark is visible in PGl (blue arrow).

**Figure 2–figure supplement 1.**
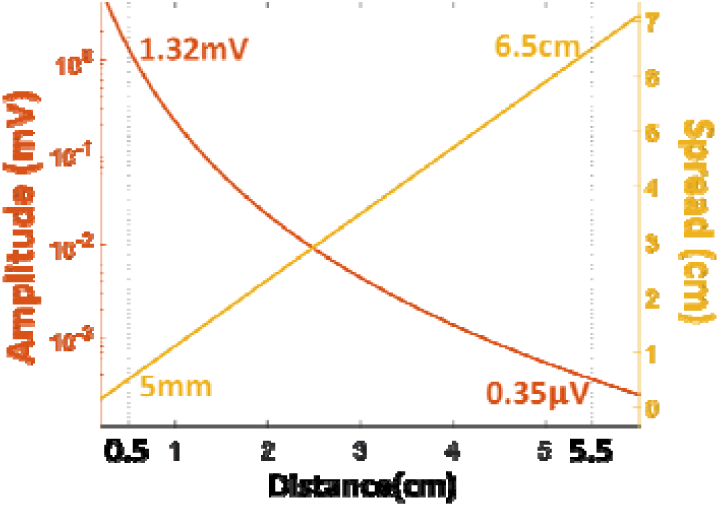
The amplitude (left ordinate, log scale) and spread (right ordinate, linear scale) of the object’s electric image on the skin, as functions of distance, based on a previously published model of electric fields in this fish (Chen et al., 2005).

**Figure 2–figure supplement 2.**
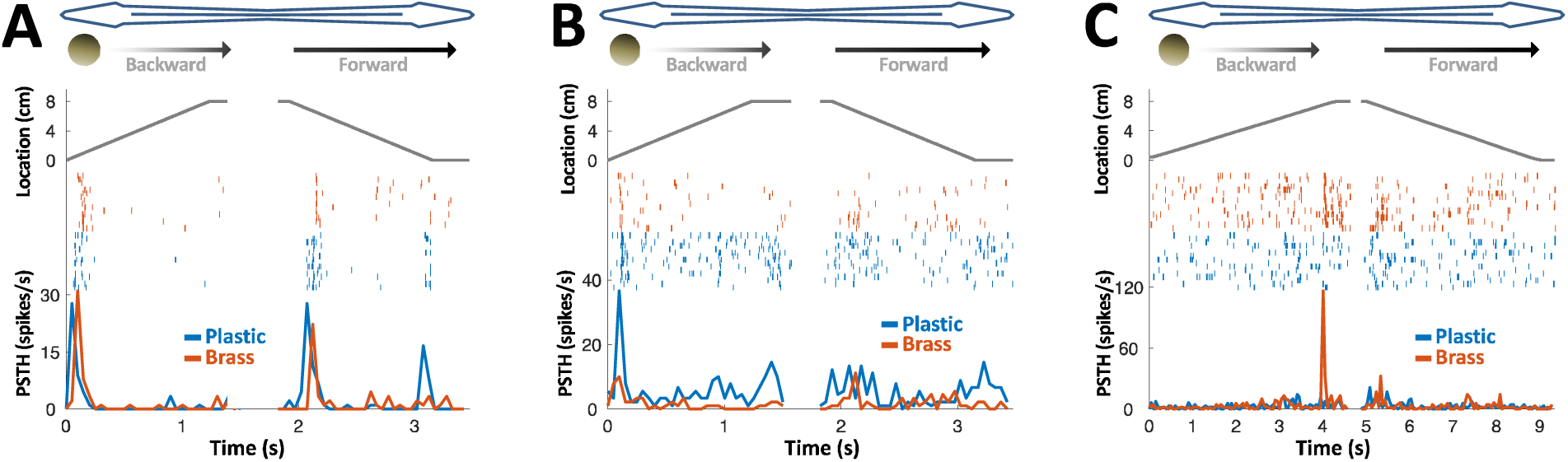
PG response to conductive/non-conductive objects. **(A-C)** Top: object location (distance from the tip of the nose) during longitudinal object motion; middle: spike raster; bottom: PSTH. Red: conductive object (brass sphere, mimicking plants and animals); Blue: non-conductive object (plastic sphere, mimicking rocks). **(A)** Example cell responding to both brass and plastic spheres (such responses were found in 57% of the cells, 24/42). **(B)** Example cell responding primarily to plastic, with a very faint response to brass (5% of the cells, 2/42). **(c)** Example cell responding to brass only (38% of the cells, 16/42). The predominance of cells responding to both brass and plastic spheres or to brass-only spheres is likely inherited from OT (Bastian, 1982).

**Figure 3–figure supplement 1.**
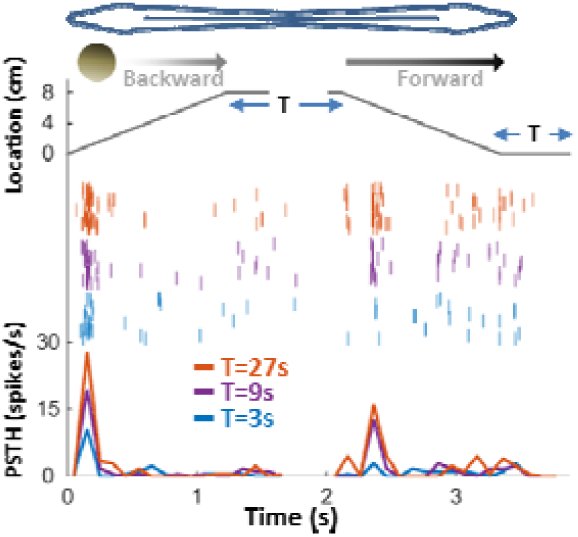
Response adaptation to periodic motion. An example motion-onset responsive cell was stimulated with repetitive back-and-forth longitudinal motion. Duration of wait intervals between movements (T) was 3, 9 or 27s (color-coded as blue, purple and red, respectively). Top: object position relative to head; middle: spike raster; bottom: PSTHs. 45% (15/33) of PG cells exhibited pronounced adaptation to repeated motion, with time-constants on the order of seconds to tens of seconds. In this example, the response strength for *T* = 27 s was larger than for *T* = 9 s, which in turn was larger than for *T* = 3 s – indicating a time-constant of adaptation >>1s.

**Figure 3–figure supplement 2.**
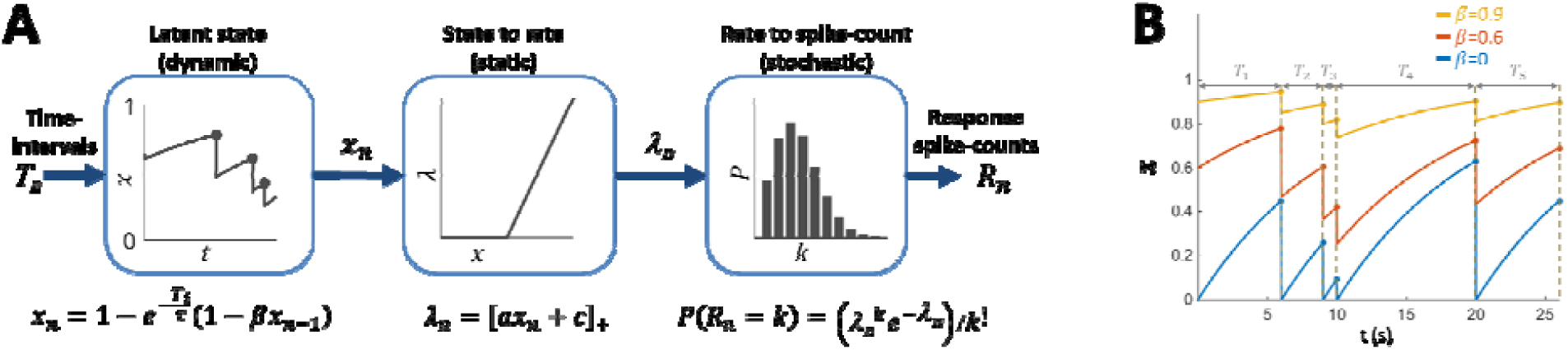
(A) Block diagram of adaptation model, which includes three stages: a dynamic latent state variable with memory parameter and recovery time-constant; a static non-negative mapping from state to firing parameter; and a stochastic spike-count generator (Poisson random variable with parameter). **(B)** Examples of state variable dynamics due to a random encounter sequence, for different values of. Note that when, each encounter entails complete reset of to 0, thus generating the history independence property.

**Figure 3–figure supplement 3.**
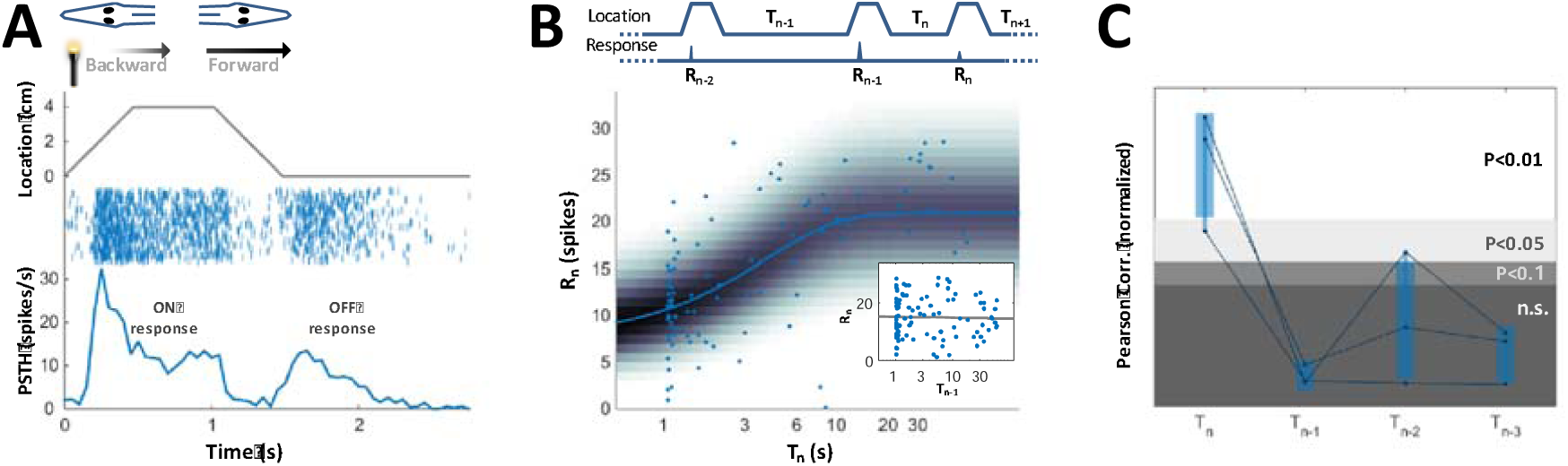
PG Visual responses. **(A)** Response of an example PG cell to longitudinal motion of a light source. Note the strong onset and offset responses. **(B)** Circles: ON response ({R_i_}, total number of spikes during backward motion) of the cell depicted in panel A, as a function of the duration of the preceding time-interval {T_i_} (1-30 s). Dynamic adaptation model was fitted to the data (β=0 for this cell); solid blue line: response rate of model fitted to the data-points; background shading: spike count distribution of the model. ON response intensity of this cell is highly correlated with the duration of the last interval (correlation=0.467, *P* < 10^−6^, random permutations). Inset: same data-points plotted as function of penultimate intervals; grey line: linear regression to log(T) (correlation= –0.022, *P* = 0.75, random permutations). **(C)** Correlations between responses and time-intervals – for the last intervals and one-, two-or three-before last intervals (*T*_*n*_, *T*_*n-1*_, *T*_*n-2*_, *T*_*n-3*_), normalized to the significance level using random permutations, and plotted here for all the adapting visual cells that we recorded (*n* = 3). Only the last interval (*T*_*n*_) was significantly correlated with the response of the cells – demonstrating that PG cells display history-independent adaptation in other modalities and not just the electrosensory one.

**Figure 4–figure supplement 1.**
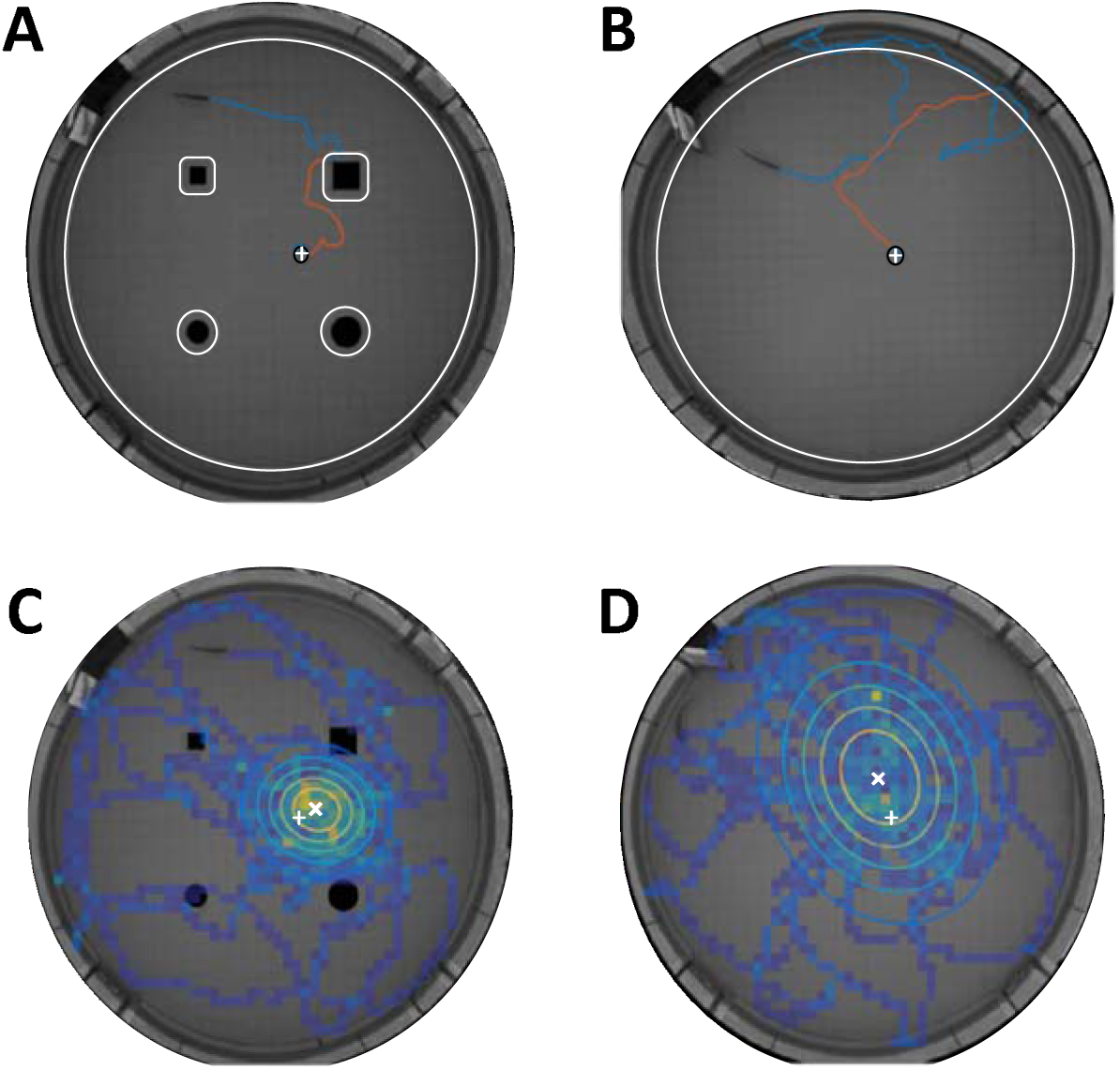
Spatial learning accuracy observed in behaving fish. As previously reported (Jun et al., 2016), fish were trained to find food in a specific location in complete darkness, either with or without the aid of landmarks. After a 12-session training stage, the animals’ performance stabilized and four test-sessions were performed. Each test session consisted of four trials, three with food present in the designated location and one ‘probe’ trial where the food was absent. The improvement in the animals’ performance (time required to find food in each trial) during the training stage (Jun et al., 2016), as well as the spatial statistics during the probe trials (see panels C, D below) indicate that the animals learnt to estimate the designated food location. Since only short range electrosensation was available to the fish, this indicates the use of path-integration from the home location, the tank walls or the encountered landmarks. For the current study, we re-analyzed the data reported in Jun et al. (2016) to produce path-integration related measures: the time/distance from last encounter to the food (A and B) and the temporal/spatial errors (C and D). **(A)** The cruising time/distance (orange line) from the last encounter (either 3 cm from landmark or 6 cm from wall, white contours) to the detection of the food (3 cm from food, black contour around white ‘+’ sign) was quantified in each of the food trials. The example shows an experiment with landmarks (black silhouettes). (**B**) Example of a food trial (similar to A) in an experiment with no landmarks. As expected, the cruising time/distance (orange) was larger in the absence of landmarks. (**C**) The spatial estimation errors were obtained by measuring where the fish searched for the missing food in the probe trials. The example depicted is of such a probe trial with landmarks (black silhouettes). Heat map: visit histogram; colored ovals: 2-D Gaussian fitting of histogram; the spatial error is the distance from fitted Gaussian center (white ‘x’) to the memorized food location (white ‘+’). (**D**) Probe trial without landmarks. Note the increase in error compared with panel C. These behavioral data enabled estimation of the number of PG cells required to achieve the temporal/spatial acuity observed (see **Figure 4**).

**Figure 4–figure supplement 2.**
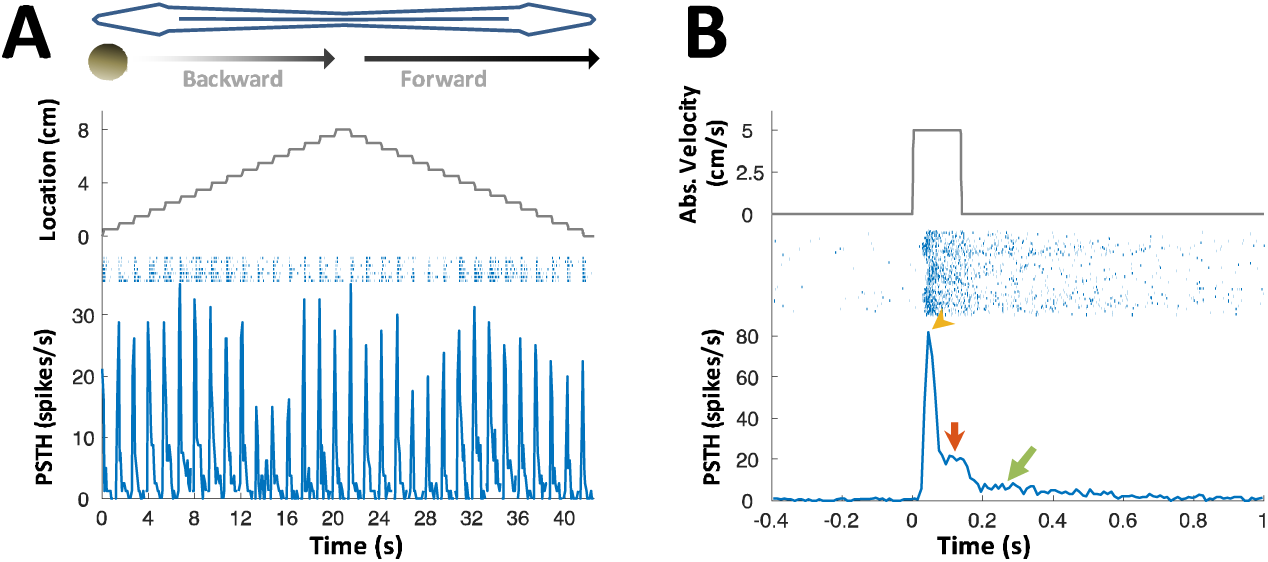
PG lateral-line responses. Response of an example PG cell to longitudinal motion of an agar sphere (15% agarose dissolved in tank water). The sphere’s conductivity was identical to that of the surrounding water, making it electrically invisible (Heiligenberg, 1973). Acoustic stimuli were also delivered to exclude the possibility of an auditory response. The relatively small number of lateral-line sensitive cells found (N=7) precludes systematic characterization of the response properties of these cells. (**A**) Motion in each direction was broken into 16 segments separated by wait period (similar to **Figure 2C**). The cell responded to this motion in either direction, anywhere across the body surface. Since the object was electrically invisible, these responses likely reflect activation of the lateral-line system, which was shown to project to PG (Giassi et al., 2012). (**B**) Cell response, averaged across body locations. Top: absolute velocity (cm/s); middle: spike raster plot; bottom: PSTH. Note that unlike most electrosensory PG cells (see **Figure 2**), this cell displayed not only motion onset response (yellow arrowhead), but also a response to continuous motion (red arrow) and prolonged response after motion cessation (green arrow; this post-stimulus discharge likely reflects wave reverberations in the experimental tank). This continuous response mode suggests that lateral-line related activity in PG conveys swimming velocity information, which can then be combined in the pallium (DL) with the time-interval information reported here to produce allocentric distance estimation between objects.

